# Developmental relationships between the human alpha rhythm and intrinsic neural timescales are dependent on neural hierarchy

**DOI:** 10.1101/2025.08.21.671637

**Authors:** Jesse T. Miles, Kurt E. Weaver, Sara Jane Webb, Jeffrey G. Ojemann

## Abstract

Maturation of human brain structure has been well-studied, but developmental changes to brain physiology are not as well understood. One consistent finding is that the peak alpha rhythm frequency (PAF) increases throughout childhood. Another is that resting-state functional connectivity shifts from sensorimotor regions in children to association regions in adolescents, a reorganization along a hierarchy called the sensorimotor-to-association (S-A) axis. In mature brains, the S-A axis has been parcellated physiologically using the duration of persistent neural activity, known as the intrinsic neural timescale (INT), which increases along the hierarchy. Here we studied the development of PAF and INT in a cohort of epilepsy patients 3 – 33 years of age undergoing intracranial electrocorticographic (ECoG) monitoring. Given the well-known developmental trajectory of PAF, and the ability to delineate hierarchy using INT, we hypothesized that changes to PAF and INT would correlate across development, but that their relationship may be influenced by hierarchy. Consequently, we predicted that age-dependent PAF increases would accompany INT decreases, and we tested whether their relationship varied between sensorimotor and association regions. We found that PAF increased significantly with age in both sensorimotor and association regions, while age-dependent INT decreases were only significant in association regions. Supporting this finding, we found a strong negative relationship between PAF and INT that was specific to association regions. Together, our results suggest that developmental divisions across the S-A axis manifest in the relationships between neurophysiological measures, providing further evidence that asynchronous development along the S-A axis depends on maturation of brain function.

**New & Noteworthy:** We report a novel developmental relationship between the human resting-state alpha rhythm frequency and the duration of intrinsic neural timescales. Using resting-state electrocorticography, we found that alpha frequency increased with age at either end of the sensorimotor-to-association cortical hierarchy, while intrinsic neural timescales only decreased with age in association regions. This negative correlation between alpha frequency and intrinsic timescale was only evident in association regions, further linking functional maturation and cortical hierarchy.

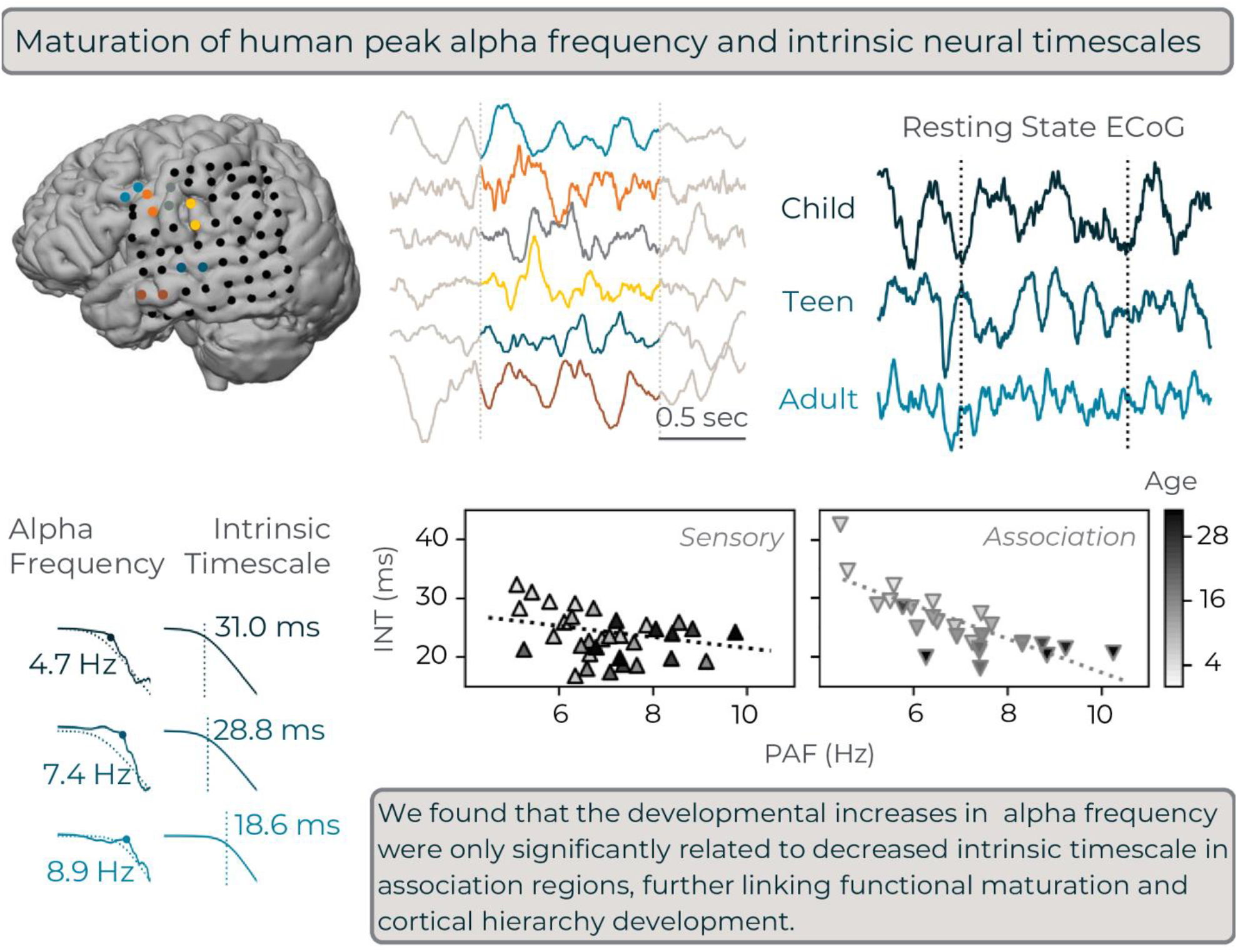

## Introduction

Human neurophysiological measures – such as electric fields recorded by electroencephalograph (EEG) – develop for years postnatally, likely mirroring the prolonged timeline of structural brain maturation (1). The peak alpha rhythm frequency (PAF), for example, increases from approximately 6 to 8 Hz in infancy and early childhood (2–4) to 8 to 12 Hz in later childhood and adolescence (5–13). In another example of prolonged functional development, neural transmission speeds increase past the age of 30 (14).

Prior evidence suggests that functional brain development occurs hierarchically. In classic models of neural processing hierarchy, information is first processed in sensorimotor regions then in association regions (15,16), forming what has become known as the sensorimotor-to-association axis (S-A axis, reviewed in 17,18). Functional maturation of the cortex appears to follow a similar progression. In children under 12, for example, fMRI-derived functional connectivity is strongest between sensory, motor, and visual cortical regions. By adolescence, however, connectivity is strongest between frontal, temporal, and parietal association regions, resembling the adult default mode network (19–22).

Hierarchy can be defined physiologically by the intrinsic timescale of neural activity. Based on temporal receptive windows (23,24), the intrinsic neural timescale (INT) quantifies the duration of a region’s correlated activity, using autocorrelation decay (23,25–32) or features of neural power spectra (33–35). In adults, sensorimotor regions tend to have shorter INTs than association regions (33,36). Hierarchies based on INT durations closely recapitulate the S-A axis regardless of whether constructed using fMRI (24,27,28), neural spike trains (25,26,30), magnetoencephalography (MEG) (29), or intracranial field potentials (23,31,33,35). Moreover, similar INT-based hierarchies emerge across mammalian species (25,28,30), suggesting INT is a robust and generalizable measure of neural processing hierarchy.

While INTs are known to increase along the S-A axis in adults, the dominant low-frequency cortical rhythm (typically in the alpha range) decreases in frequency ascending the cortical hierarchy (37). In support of more direct relationship between these measures, a scalp-EEG study in adults found PAF and INT to be negatively correlated during consciousness (38). Despite hints of physiological coupling between PAF and INT, little is known about how INTs are structured throughout the brain during childhood and adolescence. A study comparing adult and infant INTs using scalp-EEGs found that INTs tended to be longer in infants than adults (39). Similarly, infant neural event segmentation tended to be relatively uniform and coarse across the brain when measured with fMRI, whereas adults displayed hierarchical gradients of timescale segmentation (40). Timescale changes may also vary by age and region (33), but these changes have not been well studied in youths below the age of 12.

The similarities between PAF and INT prompted us to hypothesize that they would follow comparable developmental trajectories, but that those trajectories may differ based on cortical hierarchy. Namely, we predicted that the age-dependent increase in PAF would differ between sensorimotor and association regions. Further, we predicted that age-dependent PAF increases would correlate with INT decreases. Additionally, since each measure has been shown to have relationships to processing hierarchies in adults, we asked whether age-based changes to INT, and relationships between INT and PAF, were different between sensorimotor and association regions.

To test these predictions, we analyzed resting-state intracranial-EEG (iEEG) from electrocorticography (ECoG) in a cross-sectional cohort of patients, aged 3 to 33, undergoing monitoring for treatment-resistant epilepsy. We assessed hierarchy-based developmental differences by comparing PAF and INT from regions near each pole of the S-A axis (3 sensorimotor regions, and 3 association regions). Our results suggest that age-dependent PAF increases were similar in both sensorimotor and association regions, while INT decreases were only significant in association regions. This hierarchy dependence also carried over into the correlations between PAF and INT, which were significant in association regions but not sensorimotor regions.

## Methods

### Participants

Participants were originally recruited for research participation from epilepsy monitoring units at Seattle Children’s Hospital and Harborview Medical Center in Seattle, Washington. All participants underwent informed consent prior to research participation at the time of data collection and signed consent or assent forms (depending on age) granting use of their medical data for research purposes. Parental consent was also obtained for minors. Data have subsequently been approved for retrospective review by the Seattle Children’s Research Institute and University of Washington Institutional Review Boards. This study analyzes a cross-sectional cohort of participants (*n* = 13; 6 male, 7 female; Table 1) with electrocorticography grids and/or strip electrodes placed such that a mix of both sensorimotor and association regions had at least two neighboring contacts (see *Regions of interest and channel selection* for more details). All data were collected during dedicated resting-state periods, where participants were asked to quietly lay upright for at least three minutes prior to other research experiments.

**Table 1.** Patient demographics and characteristics.

| ID | Age (Y) | Sex | Electrodes | Coverage | Epileptiform Zones | Analyzed |
| --- | --- | --- | --- | --- | --- | --- |
| P1 | 3.42 | M | 8 x 8 L grid, 1 x 8 L depth (2) | Lateral peri-central and temporal | Superior post-central, inferior precentral (subclinical) | MFG, PrCG, PoCG, SMG, STG, MTG |
| P2 | 4.67 | F | 2 x 8 L strips (3), 1 x 8-16 L depth (10) | Frontal, peri-central, anterior parietal | Anterior frontal and frontal depth (subclinical) | MFG, PrCG, PoCG, SMG, STG |
| P3 | 6.67 | F | 8 x 8 L grid, 1 x 8 L strips (2) | Lateral peri-central and temporal | Peri-central discharges (subclinical) | MFG, PrCG, PoCG, SMG, STG, MTG |
| P4 | 7.25 | F | 8 x 8 R grid, 2 x 8 R strips, 1 x 8 R strip | Lateral peri-central | Superior and middle frontal | PrCG, PoCG, SMG, STG |
| P5 | 10.33 | F | 8 x 8 R grid | Lateral peri-central and temporal | Peri-central | MFG, SMG, STG, MTG |
| P6 | 10.75 | F | 8 x 8 L grid, 1 x 8-12 L strips (2) | Posterior parietal and temporal | Posterior and middle/inferior temporal | PoCG, SMG, STG, MTG |
| P7 | 13.33 | M | 8 x 8 L grid, 1 x 8 L strips (3) | Lateral peri-central and temporal | Middle/inferior temporal | MFG, PrCG, PoCG, SMG, STG, MTG |
| P8 | 15.33 | F | 8 x 8 L grid, 4 x 8 L grid, 2 x 8 strips, 1 x 8-12 strips (2), 1 x 8 depth | Peri-central to parietal cortices (posterior grid) | Posterior parietal | PrCG, PoCG, SMG |
| P9 | 15.92 | M | 8 x 8 R grid, 1 x 8 R depth (2) | Posterior frontal, lateral peri-central, parietal | Not captured | MFG, PrCG, PoCG, SMG |
| P10 | 20.75 | M | 8 x 8 L grid (split), 2 x 8 L strips, 1 x 8 L strips | Parietal and posterior temporal | Superior parietal cortex, temporal interictal spikes | PoCG, SMG, STG, MTG |
| P11 | 26.25 | M | 8 x 8 R grid | Posterior frontal, lateral peri-central, parietal | Right fronto-parietal cortex | MFG, PrCG, PoCG, SMG, STG |
| P12 | 31.33 | F | 8 x 8 R grid, 2 x 8 R strips, 1 x 6/8 R strips (4) | Lateral peri-central, temporal, parietal | Lateral, superior, posterior right temporal, mesial temporal | PrCG, PoCG, STG, MTG |
| P13 | 33.92 | M | 8 x 8 L grid, 1 x 8 strips (4) | Lateral peri-central, posterior temporal, parietal | Lateral, inferior left parietal | MFG, PrCG, PoCG, SMG, STG, MTG |

*Abbreviations – MFG, middle frontal gyrus; PrCG, precentral gyrus; PoCG, postcentral gyrus; SMG, supramarginal gyrus; STG, superior temporal gyrus; MTG, middle temporal gyrus*

### Region of interest and channel selection

Inclusion in this study required dedicated resting-state recordings, access to prior clinical notes, no clinically identified seizures during research sessions, no prior surgical resections, and ECoG coverage from a mix of sensorimotor and association region that were not in seizure onset zones. We chose to analyze regions covered in at least 8 patients to test our hypotheses about hierarchical neural development. ECoG contact locations were identified by clinical MRI reconstructions aligned to CT scans. Electrode orientations (and, by extension, channel numbering) were cross-referenced to a combination of intra-operative placement photos, surgical notes, and clinical monitoring notes.Most electrode placements covered lateral frontal, parietal, and temporal regions, so we chose a mix of cortical modalities (sensorimotor or association regions) from these areas.

With respect to cortical hierarchy – contacts covering pre-and post-central gyri were taken to represent motor and somatosensory regions, respectively, and contacts on mid-to-posterior superior temporal gyrus were considered representing early (though not primary) sensory (auditory) cortex. Collectively, these were considered sensorimotor regions. In contrast, contacts over supramarginal, middle temporal, and middle frontal gyri were chosen as association regions. Figure 1A shows an example of regions analyzed in this study in an adolescent patient.

**Figure 1.**
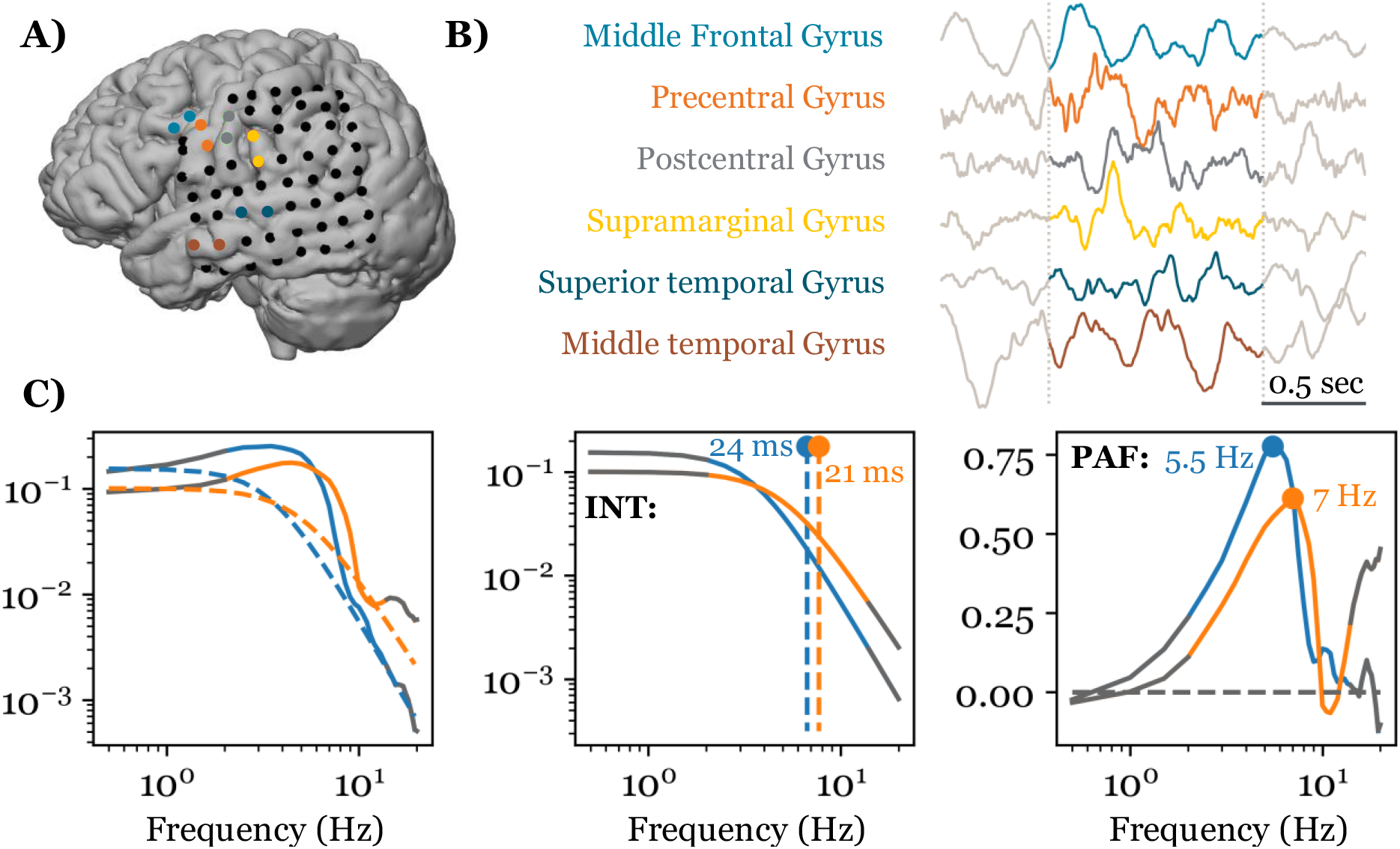
Example recording regions, iEEG data, and parameters of interest. **A)** MRI from an adolescent, black circles indicating ECoG contact locations. Colored circles indicate channels used for bipolar referencing in regions of interest. **B)** Re-referenced electrophysiology traces from each region, colored according to their grid location (labeled to the left of each trace). The segment between the vertical dashed lines is one second and traces are individually z-scored so amplitude is measured in standard deviations. **C, left)** Multitaper power spectral density estimates from the middle frontal gyrus (blue) and precentral gyrus (orange), with the frequency range of interest (3 – 14 Hz) shown in color. Estimates of the aperiodic component of the spectrum are shown as dashed, colored lines for each spectrum. **C, middle)** Model fits shown on their own, with vertical dashed lines indicating the accompanying knee frequency (*k*_*f*_) estimates used to calculate INT. **C, right)** Residual power estimates after correcting for aperiodic activity with circles overlying each spectrum’s peak alpha frequency.

We used clinical notes to exclude channels found to be in the seizure onset zone or that were mentioned as having frequent seizure spreading or interictal spiking activity, though no clinical seizures occurred during research sessions. Channels near regions that were suspected of having frequent interictal activity on manual inspection were also excluded, even if not mentioned clinically. Of the remaining channels, we sought two adjacent contacts within the same gyrus that could be used for bipolar referencing. When more than two remaining adjacent contacts were available, we chose pairs of contacts with the fewest identified noise segments and similar amplitude distributions. When possible, we tried to pick pairs in qualitatively comparable portions of the gyri across patients (*e*.*g*., portions of the superior temporal gyrus directly inferior to pre- or post-central gyri; segments of pre- and post-central gyri on the lateral surface, but not immediately adjacent to the Sylvian fissure; the most anterior pair of supramarginal contacts). Gyri were identified based on automated registration of Harvard-Oxford atlas labels to patient clinical neuroimaging using the Localizing Electrodes GUI (41) and manual inspection of co-registered CT and T1-weighted MR images.

### Signal processing

Signals were sampled at a minimum of 512 Hz and maximum of 4800 Hz (depending on acquisition system and site), with majority of sampling rates in the 1000 to 2000 Hz range. As such, signals were resampled to a common sampling rate of 1024 Hz using the following procedure. All signals were up-sampled by interpolation with a cubic spline to the next power of 2 greater than their original sampling rate, lowpass filtered with a 6^th^ order Butterworth filter at the Nyquist frequency of the common resampling rate (512 Hz) to prevent aliasing, and down-sampled to the final 1024 Hz sampling rate.

Electrode drift was removed by subtracting a 1 second moving average from each resampled trace. Line noise was filtered with 6^th^ order Butterworth filters at 180 Hz, 120 Hz, and 60 Hz. Before the final 60 Hz filter, we also applied a 6^th^ order low-pass Butterworth filter at 150 Hz. All re-referenced and filtered signals were screened for artifacts and possible epileptiform activity by excluding data segments with 8 or more data points above 10 standard deviations in a 0.125 second window. These events were exceedingly rare, comprising less than 0.1% of data for any given channel, and upon manual inspection were far more likely due to patient movement or adjusting electrode cables than pathological activity. Artifact-free signal segments were then z-score normalized to control for baseline amplitude differences due to variability in equipment and/or surgical placements. Figure 1B shows an example of the electrophysiological signals obtained during the resting-state after re-referencing.

### Spectral calculations

We used the pymultitaper package to calculate multi-taper spectral estimates on normalized signals. All spectral estimates were calculated as time frequency decompositions using a time-bandwidth product of 3 with 5 tapers on 1 second segments of data with 75% overlap (*i*.*e*., slid by 0.25 seconds per segment). Segments were zero padded to bring the frequency resolution to 0.25 Hz. Between three and ten minutes of resting-state data were collected for each patient, enabling creation of distributions for PAF and INT in each region. Spectral estimates for a subset of data were cross-checked against the widely used Chronux package (42) and found to be consistent.

### Spectral parameterization

Individual power spectra were parameterized following methods of (43). The decay in field potential power in both scalp and intracranial-EEG have been shown to follow a power law, meaning higher frequencies contain exponentially less power than lower frequencies (43–48). In logarithmic space (for both frequency and power), this relationship sometimes approximates a line with negative slope and y-intercept offset. In human intracranial recordings there is often a characteristic “knee” shape in the lower frequency bands, after which there is a considerably more linear relationship between power and frequency (again, in log-log space) (33,34,43,44).

The knee parameter has been proposed as a physiological measure of cortical hierarchy, due to its relationship to intraregional autocorrelation decay and temporal receptive windows (23,24,33,34). To estimate this value, we parameterized all power spectra using a modified version of the specparam software package (34,43). Briefly, the algorithm aims to parameterize periodic and aperiodic (1/f) spectral components by first fitting and subtracting the 1/*f*^*x*^ component of the spectrum, iteratively fitting and subtracting residual power spectral peaks with Gaussians, and then re-fitting the 1/*f*^*x*^ component of the spectrum after residual (peak) power has been subtracted. This process yields several parameters of interest, such as the 1/*f*^*x*^ exponent (*x*), offset, and knee frequency values of the spectrum. These parameters define the function that can be subtracted from spectra to generate 1/*f-*adjusted periodic power estimates.

In the modified fitting procedure (34), the aperiodic model is a generalization of the Lorentzian function:

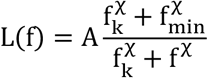

With this formulation, *A* – the power fitted at the lowest frequency analyzed (*f*_*min*_) – is the offset, *f*_*k*_ is the knee frequency, and *χ* defines the slope of power decay at frequencies higher than *f*_*k*_. The INT is calculated from *f*_*k*_ as *τ = (2πf*_*k*_)^*-1*^ (33). Notably, this formulation also fits signals that are essentially log-linear across the entire frequency range analyzed (*i*.*e*., in spectra without a knee), meaning separate models do not need to be used for fitting spectra with different shapes (34). A detailed comparison of this model and the original specparam models can be found in the supplemental materials of (34). Figure 1C shows examples of the power spectra, fitting procedure, and individual PAF and INT measurements.

To mitigate bias toward high-frequency error minimization in the fitting procedure, we also linearly interpolated frequencies between 0.5 and 1 Hz to have the same number of points as frequencies between 10 and 50 Hz, and frequencies between 1 and 10 Hz to have the same number of points as frequencies above 10 Hz prior to running the fitting routines. Upon manual inspection, this appeared to provide better fits, particularly in the low-frequency portion of the spectrum, when returning fitted values at the original frequency resolution, but we did not test the goodness of fit explicitly. We used the following hyperparameters for fitting power spectra:

1. peak_width_limits = (1, 40)
2. max_n_peaks = 6
3. aperiodic_mode = ‘lorentzian’
4. min_peak_height = 0.15
5. peak_threshold = 2

Adjusted power spectra were used to identify PAF by searching for local maxima between 3 and 14 Hz. If there were multiple peaks in the range, the largest peak was chosen. A threshold was used to exclude noisy model fits, and the base width of the peaks was required to be at least 1.5 Hz but less than 10 Hz.

### Statistical modeling

All regressions were run using linear ordinary least squares models with version 0.14.2 of the Python statsmodels package (49). Distributions of estimates for both PAF and INT were generated from time-frequency decomposition of the resting-state data (as detailed in *Spectral calculations*), and averages of these distributions were used for statistics. Depending on the analysis, averaging occurred either across time for a single region, grouped by whether regions were sensorimotor or association, or across all datapoints (regions and timepoints) per individual. When age was used as an independent variable it was log_2_-transformed to linearize its relationship with dependent variables. Exploratory regressions that split regions by hierarchy membership were considered significant for *p-*values below 0.025 (the Bonferroni-adjusted value for two comparisons). When splitting by region, we implemented the default Benjamini-Hochberg *p-*value correction (50) from statsmodels.stats.multitest.fdrcorrection to control the false discovery rate.

For comparing PAF and INT across all regions, we included age as a regressor and allowed cortical hierarchy (sensorimotor or association categorical labels) to interact with PAF in the model. This ensured our results were not simply due to collinearity with age and tested whether the linear relationship between intrinsic timescale and PAF differed depending on whether a region was sensorimotor or association. In Patsy notation for the formula API in statsmodels:

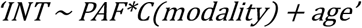

where *C* denotes use of a categorical variable. Note that statsmodels automatically includes main effect terms when looking at interactions, so they do not need to be included as separate formula entries.

## Results

### Peak alpha frequency by age and region

Averaging PAF estimates across all regions of interest per participant, we replicated the general age-dependent increase in PAF (Figure 2A, *left*; *R*^*2*^ = 0.720; *T* = 5.31; *n* = 13; *p* = 2.45 × 10^−4^). Regions were then separated based on their common designation as sensorimotor or association to test whether PAF developed differently at opposing poles of the S-A axis. Separate regressions for both sensorimotor and association regions were both significant, suggesting no clear hierarchy dependence in PAF development (Figure 2A, *center*; sensorimotor – *R*^*2*^ = 0.536; *T* = 3.57; *n* = 13; *p* = 4.43 × 10^−3^; Figure 2A, *left;* association – *R*^*2*^ = 0.733; *T* = 5.50; *n* = 13; *p* = 1.86 × 10^−4^). Regressions in individual regions were significant after false discovery rate correction for all but middle frontal and postcentral gyri (Figure 2C, Table 2).

**Table 2.** PAF regression results for individual regions.

| **Table 2 – PAF regression results for individual regions** |  |  |  |  |  |
| --- | --- | --- | --- | --- | --- |
| *Region* | *n* | *R<sup>2</sup>* | *T* | *p* | *FDR<sub>p</sub>* |
| Middle frontal gyrus | 8 | 0.098 | 0.81 | 0.4507 | 0.4507 |
| **Precentral gyrus** | **10** | **0.698** | **4.30** | **0.0026** | **0.0078\*** |
| Postcentral gyrus | 12 | 0.224 | 1.70 | 0.1203 | 0.1444 |
| **Supramarginal gyrus** | **12** | **0.701** | **4.85** | **0.0007** | **0.0041\*** |
| **Superior temporal gyrus** | **11** | **0.492** | **2.95** | **0.0161** | **0.0242\*** |
| **Middle temporal gyrus** | **8** | **0.665** | **3.45** | **0.0136** | **0.0242\*** |
| ***Bolded \* indicates significant relationship after false discovery rate correction*** |  |  |  |  |  |

**Figure 2.**
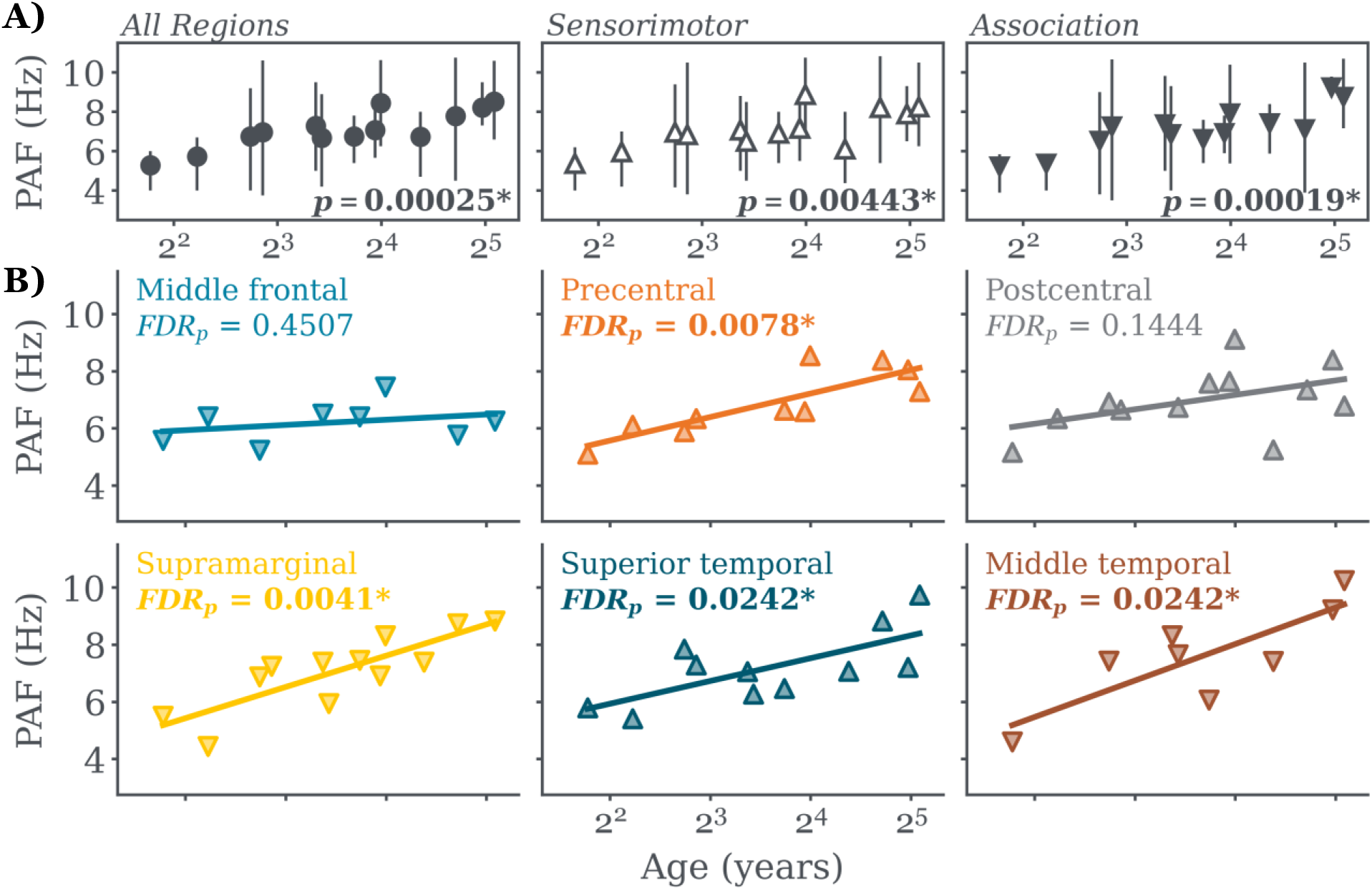
Peak alpha frequency increases with age. **A), *left*** Regression analysis comparing combined PAF from all regions per patient (*n* = 13) with age. Average pooled PAF for each patient is a circle and error bars demarcate the 25^th^ and 75^th^ percentiles of PAF values. Average PAF is significantly correlated with age. **A), *center, right*** Same as **A), *left*** but for sensorimotor and association regions, respectively. Mean PAF for sensorimotor cortex (*center*) is represented by upward, white triangles. Association cortex (*right*) is in grey, with means plotted as downward triangles. All x-axes are shown as log_2_ of age. Annotations show *p*-value for ordinary least squares regression fit. **C)** Regressions split by region. Note that the number of patients with coverage varies per region (see Table 2 for full regression details).

### Intrinsic neural timescale by age and region

When all regions per patient were pooled to estimate INT, we found a significant decrease in INT with increasing age (Figure 3A, *left*; *R*^*2*^ = 0.497; *t* = −3.30; *n* = 13; *p* = 0.0071). Regressing sensorimotor regions separately from association regions resulted in an age-dependent timescale decrease that did not withstand multiple comparison correction (critical value halved for comparison with association regions; Figure 3A *center*; sensorimotor – *R*^*2*^ = 0.354; *T* = −2.54; *n* = 13; *p* = 0.032). Association regions, however, did exhibit a significant age-dependent decrease in INT (Figure 3A, *right*; association – *R*^*2*^ = 0.608; *T* = −4.13; *n* = 13; *p* = 0.0017). Further separation into individual regions revealed all individual association regions – but no sensorimotor regions – had significant age-dependent decreases to intrinsic neural timescale that survived false discovery rate correction (Figure 3B, Table 3).

**Table 3.** INT regression results for individual regions.

| Table 3 – INT regression results for individual regions |  |  |  |  |  |
| --- | --- | --- | --- | --- | --- |
| Region | $n$ | $R^2$ | $T$ | $p$ | $FDR_p$ |
| **Middle frontal gyrus** | **8** | **0.625** | **-3.16** | **0.0195** | **0.0390\*** |
| Precentral gyrus | 10 | 0.236 | -1.57 | 0.1542 | 0.1542 |
| Postcentral gyrus | 12 | 0.361 | -2.38 | 0.0388 | 0.0582 |
| **Supramarginal gyrus** | **12** | **0.443** | **-2.82** | **0.0182** | **0.0390\*** |
| Superior temporal gyrus | 11 | 0.286 | -1.89 | 0.0901 | 0.1081 |
| **Middle temporal gyrus** | **8** | **0.783** | **-4.65** | **0.0035** | **0.0209\*** |
| **Bolded \*** indicates significant relationship after false discovery rate correction |  |  |  |  |  |

**Figure 3.**
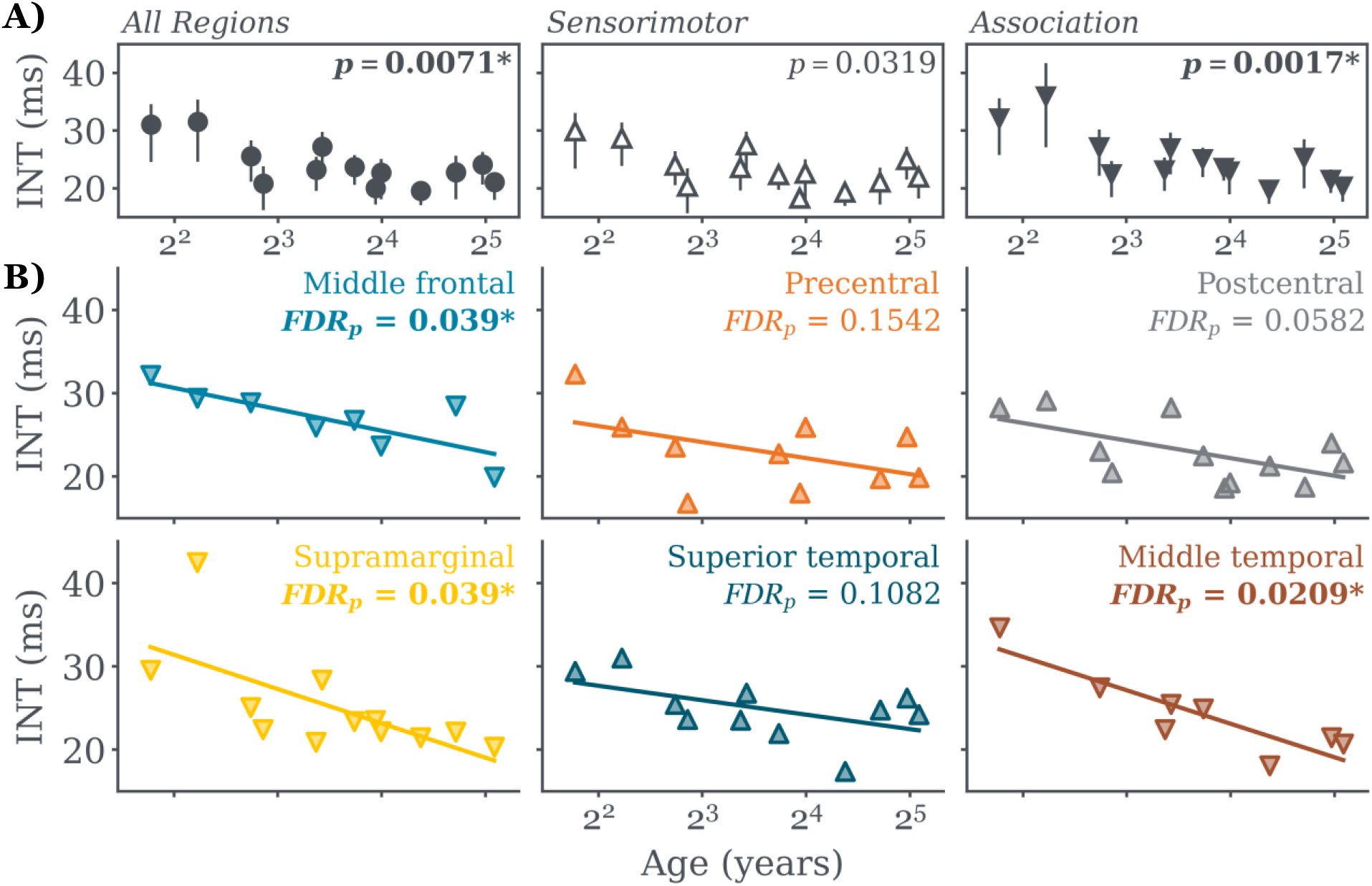
Hierarchical age-dependence of intrinsic neural timescale. **A) *left***, INT is negatively correlated with age when combining all regions per patient. **A) *center, right***, as in **A) *left***, but with regions grouped by hierarchy. A negative relationship between INT and age is only significant in association cortex after multiple comparison correction. **B)** When regressed individually, all association regions, but no sensori-motor regions, exhibit significant age-dependent decreases in INT, but no relationships between age and INT after false discovery rate correction. All X-axes are shown as log_2_ of age

### Relation between PAF, INT, and S-A hierarchy

To test our prediction that INT and PAF would have a significant negative relationship, we correlated the average regional PAF against the average regional INT for all patients in the dataset. We found a significant correlation between PAF and INT when combining data across regions and ages (Figure 4A, Pearson’s *R* = 0. 573, *n* = 61 regions; *p* = 1.369 × 10^−6^, not adjusted for age-dependence).

**Figure 4.**
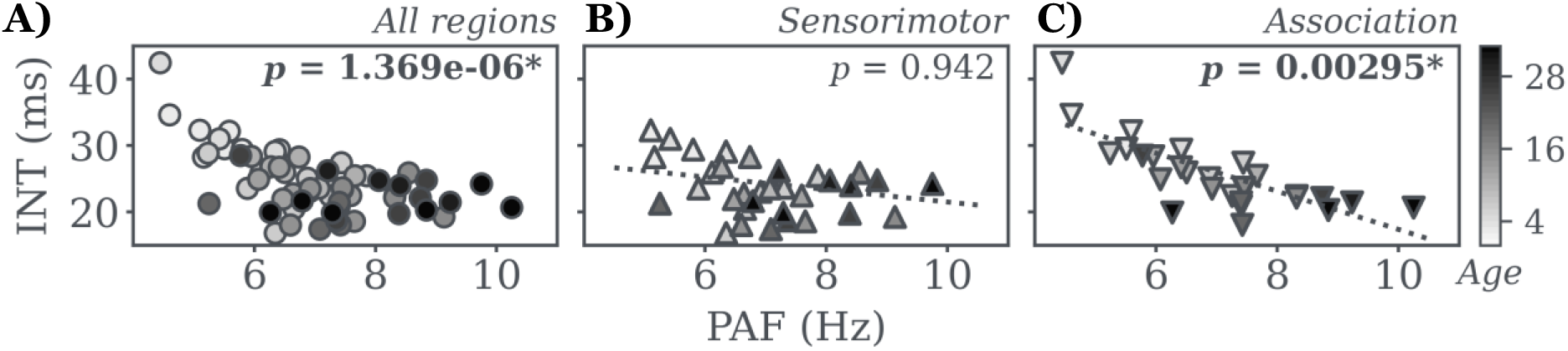
Peak alpha frequency and intrinsic neural timescale are correlated. **A)** INT is negatively related to PAF. **B)** The relationship between INT and PAF is not significant in sensorimotor regions. **C)** The relationship between INT and PAF in association regions remains significant after separating from sensorimotor regions and accounting for age-dependence.

Because we found evidence for hierarchically dependent changes in INT (Figure 3), we performed a follow-up multiple linear regression analysis that allowed PAF to interact with cortical hierarchy (sensorimotor vs. association), and, since both measures were significantly associated with age, we also added age as a regressor. Consistent with hierarchical differences in the relationship between INT and PAF, we found a significant categorical main effect of the cortical modality term of the model, as well as a significant interaction between PAF and cortical modality. As before (Figure 3), the relationship between INT and age also remained significant in the multiple regression model (Figure 4A; full model adjusted *R*^*2*^ = 0.497; *F* = 15.83; *n* = 61; *p*_*mod*_ = 0.0027, *p*_*PAF*_ = 0.0008, *p*_*mod:PAF*_ = 0.0064, *p*_*age*_ = 0.0005).

A confirmatory *post hoc* analysis revealed that while sensorimotor regions did not exhibit significant relationships between INT and PAF, (Figure 4B; sensorimotor – adjusted *R*^*2*^ = 0.231, *F* = 5.80, *n* = 33, *p*_*PAF*_ = 0.9423, *p*_*age*_ = 0.0138), association regions did have a significant relationship between INT and PAF (Figure 4C; association – adjusted *R*^*2*^ = 0.649, *F* = 25.97, *n* = 38, *p*_*PAF*_ = 0.0030, *p*_*age*_ = 0.0173). Together, these findings suggest that the negative relationship between INT and PAF is specific to association regions and is not solely dependent on age.

## Discussion

Results from this study partially supported our hypothesis that PAF and INT would follow similar, hierarchy-dependent developmental trajectories. We replicated the expected age-dependent increase in PAF using intracranial recordings, but did not find evidence that PAF development was linked to hierarchy (Figure 2). As predicted, INT generally decreased with age (Figure 3A), but we only found evidence for a significant age-dependent INT decrease in association regions (Figure 3B). At the regional level, none of the sensorimotor regions included in the study had significant age-dependent INT decreases, while all association regions did. There is some suggestion that sensorimotor regions may have decreased to a stable minimum INT at an earlier age, but a larger sample and age-range would be needed to confirm this speculation. Of note, we found a significant relationship between PAF and INT, such that increased PAF was associated with decreased INT (Figure 4A). Further, the negative relationship between INT and PAF was only significant in association regions (Figure 4B, C), mirroring results for INT age-dependence.

Much like our findings of age-dependent PAF increases, Johnson and colleagues (2022) used iEEG to report a pair of peaks in fronto-temporal rhythmic activity that also undergo age-related frequency changes in a sample of 6-to-21-year-olds completing a working memory task. Specifically, the lower frequency peak (approx. 2 – 4 Hz; “slow theta”) decreased in frequency with age, while a higher frequency peak (approx. 6-8 Hz; “fast theta”) increased with age (51). Given the similarities between those results and ours, it is unclear whether development of the “fast theta” rhythm is distinct from that of PAF seen here – especially as different regions are known to exhibit different spectral characteristics (52,53), and these data were collected during the resting-state.

In addition, a large iEEG study found that the magnitude of the frequency-dependent decay in spectral power (the spectral slope) changed across development, with age-dependent changes that varied by attentional state in some regions (54). Interestingly, comparisons of cortical thickness with spectral slope suggested that different regions may have opposing relationships, precluding a simple explanation of cortical thickness as a developmental mediator of spectral slope. Regional peak rhythm frequencies in adults, on the other hand, are known to vary with cortical thickness (37), as are INTs (33). A robust literature describing spectral slope decreases with development has emerged in scalp EEG studies (*e*.*g*., Caffarra et al. 2024; Cellier et al. 2021; Tröndle et al. 2022), but further studies that can reliably pinpoint regional developmental differences of both periodic and aperiodic activity are necessary to understand why these changes emerge, and whether any patterns govern their variation across brain regions.

When considering age-dependent changes to neurophysiology, age is generally a proxy for structural changes that occur during brain maturation. For example, intracortical myelin content is increased near the sensorimotor pole of the S-A axis in adults (33,55,56), matures in a gradient along the S-A axis (22,55,57,58), and exhibits weaker age-related changes in regions near the association pole (55,59). The S-A axis is also recapitulated in cortical thickness measurements; sensory regions are thinner than association regions in mature brains (60,61) and developmental trajectories of cortical thickness tend to differ across the S-A axis (62–64). Some of these structural measures may inform the hierarchy-based differences we found, such as the differences in INT development for sensorimotor and association regions. An important future direction will be assessing whether structural measures help explain the hierarchy-dependent relationship between PAF and INT.

Due to its basis in correlated activity and temporal receptive windows, INT has been conceptualized as a measure of neural integration (reviewed in 65). In this perspective, regions with short INTs process information quickly, while longer INTs allow multiple information streams to be combined into abstract representations (23–25,31,33). This aligns with hierarchy-based INT tiling, where shorter INTs are seen toward the sensory pole of the S-A axis. Alpha, on the other hand, is often considered an inhibitory rhythm, involved in sensory perception and attention (66–68). PAF, specifically, has been linked to perceptual change processing (69,70). These varied observations suggest that alpha may serve multiple roles: a network-wide index of inhibition in regions with correlated alpha power increases, and an attention spotlight in regions with phase-aligned PAF changes. Together, INT and PAF may form complimentary components of a dual-process, whereby phase-aligned PAF enables fast information sharing between regions during selective attention and INT sets the duration over which information streams can be combined. As such, coordination between the two could regulate proper information segregation (mediated by alpha power co-increases), transmission (mediated by phase-aligned PAF), and integration (mediated by INT) as processing proceeds through cortical hierarchies.

Though speculative, such an organization may help explain why we found a significant relationship between PAF and INT in association, but not sensorimotor, regions. Multi-sensory integrative processing has limited influence in sensorimotor regions, where alpha is more likely involved in sampling external, sensory information, and seems to play a generally inhibitory role. In regions farther up the hierarchy, however, alpha may be related to sampling from internal sources of information, as would be expected for selective attention. Such a model aligns with observations of differential alpha generation across the cortical hierarchy (71,72), potentially lending to distinct roles for alpha, and with findings that INTs are modulated by allocation of sensory attention (73).

We based our initial hypothesis on studies of both intracranial (human and model organism) and noninvasive neurophysiology data. Accordingly, it is unlikely that our predictions would have held if our results were driven by epilepsy-related physiology. Although it is plausible that epilepsy could alter network-level phenomena, like cortical hierarchy formation, the more parsimonious interpretation is that our findings are applicable to non-pathological brain function. Still, testing the reliability of our results using non-invasive protocols in normative control populations (*e*.*g*., with MEG) and in model systems will be important for assessing the generalizability of our observations.

Our results establish developmental relationships between two physiological markers of information processing, PAF and INT, using resting-state iEEG data across a broad age-range. Alpha frequency increases occurred across the cortical hierarchy, while development of INT varied depending on hierarchy rank. Similarly, the relationship between PAF and INT was clear in association regions, but absent in sensorimotor regions. We hope our finding of a strong, hierarchy-dependent link between these measures spurs investigation into their mechanistic relationship and examination of potential alterations to their relationship in atypical neural function.

## Acknowledgements

We are grateful to the patients who dedicated their time to participating in research, and to the Harborview and Seattle Children’s Hospital staff for their help facilitating our work and caring for patients. Many thanks to members of the Grid Lab at the University of Washington and the PBS lab at Seattle Children’s Research Institute for helpful comments on earlier drafts of the manuscript.

## Author Contributions

JTM, KEW, SJW, and JGO conceived and designed research. JTM analyzed data. KEW performed experiments. JTM, KEW, SJW, and JGO interpreted results. JTM prepared figures and drafted the manuscript. JTM, KEW, SJW, and JGO edited, revised, and approved the final version of the manuscript.

## Disclosures

The authors have no conflicts of interest to disclose.

## Funding

This work was made possible by funds from the Richard G. Ellenbogen Endowed Chair for Pediatric Neurosurgery (JGO).

## Data and code availability

Code and anonymized data will be made publicly available after peer review, or upon request to the authors

## Notes

### Competing Interest Statement

The authors have declared no competing interest.

## References

1. Bethlehem R a. I, Seidlitz J, White SR, Vogel JW, Anderson KM, Adamson C, et al. Brain charts for the human lifespan. Nature. 2022 Apr;604(7906):525–33.

2. Marshall PJ, Bar-Haim Y, Fox NA. Development of the EEG from 5 months to 4 years of age. Clinical Neurophysiology. 2002 Aug;113(8):1199–208.

3. Berchicci M, Zhang T, Romero L, Peters A, Annett R, Teuscher U, et al. Development of Mu Rhythm in Infants and Preschool Children. Dev Neurosci. 2011;33(2):130–43.

4. Schaworonkow N, Voytek B. Longitudinal changes in aperiodic and periodic activity in electrophysiological recordings in the first seven months of life. Developmental Cognitive Neuroscience. 2021 Feb 1;47:100895.

5. Miskovic V, Ma X, Chou CA, Fan M, Owens M, Sayama H, et al. Developmental changes in spontaneous electrocortical activity and network organization from early to late childhood. NeuroImage. 2015 Sep;118:237–47.

6. Thorpe SG, Cannon EN, Fox NA. Spectral and source structural development of mu and alpha rhythms from infancy through adulthood. Clinical Neurophysiology. 2016 Jan;127(1):254–69.

7. Cellier D, Riddle J, Petersen I, Hwang K. The development of theta and alpha neural oscillations from ages 3 to 24 years. Developmental Cognitive Neuroscience. 2021 Aug 1;50:100969.

8. Candelaria-Cook FT, Schendel ME, Flynn L, Cerros C, Kodituwakku P, Bakhireva LN, et al. Decreased resting-state alpha peak frequency in children and adolescents with fetal alcohol spectrum disorders or prenatal alcohol exposure. Developmental Cognitive Neuroscience. 2022 Oct;57:101137.

9. Freschl J, Azizi LA, Balboa L, Kaldy Z, Blaser E. The development of peak alpha frequency from infancy to adolescence and its role in visual temporal processing: A meta-analysis. Developmental Cognitive Neuroscience. 2022 Oct;57:101146.

10. Tröndle M, Popov T, Dziemian S, Langer N. Decomposing the role of alpha oscillations during brain maturation. eLife. 2022 Aug 25;11:e77571.

11. Caffarra S, Kanopka K, Kruper J, Richie-Halford A, Roy E, Rokem A, et al. Development of the Alpha Rhythm Is Linked to Visual White Matter Pathways and Visual Detection Performance. J Neurosci. 2024 Feb 7;44(6):e0684232023.

12. Vandewouw MM, Sato J, Safar K, Rhodes N, Taylor MJ. The development of aperiodic and periodic resting-state power between early childhood and adulthood: New insights from optically pumped magnetometers. Developmental Cognitive Neuroscience. 2024 Oct 1;69:101433.

13. Hill AT, Clark GM, Bigelow FJ, Lum JAG, Enticott PG. Periodic and aperiodic neural activity displays age-dependent changes across early-to-middle childhood. Developmental Cognitive Neuroscience. 2022 Apr;54:101076.

14. Van Blooijs D, Van Den Boom MA, Van Der Aar JF, Huiskamp GM, Castegnaro G, Demuru M, et al. Developmental trajectory of transmission speed in the human brain. Nat Neurosci. 2023 Apr;26(4):537–41.

15. Mesulam MM. From sensation to cognition. Brain. 1998 Jun 1;121(6):1013–52.

16. Felleman DJ, Van Essen DC. Distributed hierarchical processing in the primate cerebral cortex. Cereb Cortex. 1991;1(1):1–47.

17. Huntenburg JM, Bazin PL, Margulies DS. Large-Scale Gradients in Human Cortical Organization. Trends in Cognitive Sciences. 2018 Jan 1;22(1):21–31.

18. Sydnor VJ, Larsen B, Bassett DS, Alexander-Bloch A, Fair DA, Liston C, et al. Neurodevelopment of the association cortices: Patterns, mechanisms, and implications for psychopathology. Neuron. 2021 Sep 15;109(18):2820–46.

19. Dong HM, Margulies DS, Zuo XN, Holmes AJ. Shifting gradients of macroscale cortical organization mark the transition from childhood to adolescence. Proc Natl Acad Sci USA. 2021 Jul 13;118(28):e2024448118.

20. Luo AC, Sydnor VJ, Pines A, Larsen B, Alexander-Bloch AF, Cieslak M, et al. Functional connectivity development along the sensorimotor-association axis enhances the cortical hierarchy. Nat Commun. 2024 Apr 25;15(1):3511.

21. Margulies DS, Ghosh SS, Goulas A, Falkiewicz M, Huntenburg JM, Langs G, et al. Situating the default-mode network along a principal gradient of macroscale cortical organization. Proceedings of the National Academy of Sciences. 2016 Nov;113(44):12574–9.

22. Sydnor VJ, Larsen B, Seidlitz J, Adebimpe A, Alexander-Bloch AF, Bassett DS, et al. Intrinsic activity development unfolds along a sensorimotor–association cortical axis in youth. Nat Neurosci. 2023 Apr;26(4):638–49.

23. Honey CJ, Thesen T, Donner TH, Silbert LJ, Carlson CE, Devinsky O, et al. Slow Cortical Dynamics and the Accumulation of Information over Long Timescales. Neuron. 2012 Oct;76(2):423–34.

24. Hasson U, Yang E, Vallines I, Heeger DJ, Rubin N. A Hierarchy of Temporal Receptive Windows in Human Cortex. J Neurosci. 2008 Mar 5;28(10):2539–50.

25. Murray JD, Bernacchia A, Freedman DJ, Romo R, Wallis JD, Cai X, et al. A hierarchy of intrinsic timescales across primate cortex. Nat Neurosci. 2014 Dec;17(12):1661–3.

26. Wasmuht DF, Spaak E, Buschman TJ, Miller EK, Stokes MG. Intrinsic neuronal dynamics predict distinct functional roles during working memory. Nat Commun. 2018 Aug 29;9(1):3499.

27. Ito T, Hearne LJ, Cole MW. A cortical hierarchy of localized and distributed processes revealed via dissociation of task activations, connectivity changes, and intrinsic timescales. NeuroImage. 2020 Nov 1;221:117141.

28. Raut RV, Snyder AZ, Raichle ME. Hierarchical dynamics as a macroscopic organizing principle of the human brain. Proc Natl Acad Sci USA. 2020 Aug 25;117(34):20890–7.

29. Golesorkhi M, Gomez-Pilar J, Tumati S, Fraser M, Northoff G. Temporal hierarchy of intrinsic neural timescales converges with spatial core-periphery organization. Commun Biol. 2021 Mar 4;4(1):1–14.

30. Siegle JH, Jia X, Durand S, Gale S, Bennett C, Graddis N, et al. Survey of spiking in the mouse visual system reveals functional hierarchy. Nature. 2021 Apr 1;592(7852):86–92.

31. Cusinato R, Alnes SL, Maren E van, Boccalaro I, Ledergerber D, Adamantidis A, et al. Intrinsic Neural Timescales in the Temporal Lobe Support an Auditory Processing Hierarchy. J Neurosci. 2023 May 17;43(20):3696–707.

32. Song M, Shin EJ, Seo H, Soltani A, Steinmetz NA, Lee D, et al. Hierarchical gradients of multiple timescales in the mammalian forebrain. Proceedings of the National Academy of Sciences. 2024 Dec 17;121(51):e2415695121.

33. Gao R, van den Brink RL, Pfeffer T, Voytek B. Neuronal timescales are functionally dynamic and shaped by cortical microarchitecture. Vinck M, Colgin LL, Womelsdorf T, editors. eLife. 2020 Nov 23;9:e61277.

34. Bush A, Zou JF, Lipski WJ, Kokkinos V, Richardson RM. Aperiodic components of local field potentials reflect inherent differences between cortical and subcortical activity. Cerebral Cortex. 2024 May 1;34(5):bhae186.

35. Manea AMG, Maisson DJN, Voloh B, Zilverstand A, Hayden B, Zimmermann J. Neural timescales reflect behavioral demands in freely moving rhesus macaques. Nat Commun. 2024 Mar 9;15(1):2151.

36. Cusinato R, Seiler A, Schindler K, Tzovara A. Sleep Modulates Neural Timescales and Spatiotemporal Integration in the Human Cortex. J Neurosci [Internet]. 2025 Apr 9 [cited 2025 Jul 29];45(15). Available from: https://www.jneurosci.org/content/45/15/e1845242025

37. Mahjoory K, Schoffelen JM, Keitel A, Gross J. The frequency gradient of human resting-state brain oscillations follows cortical hierarchies. Dugué L, Colgin LL, Dugué L, editors. eLife. 2020 Aug 21;9:e53715.

38. Buccellato A, Zang D, Zilio F, Gomez-Pilar J, Wang Z, Qi Z, et al. Disrupted relationship between intrinsic neural timescales and alpha peak frequency during unconscious states – A high-density EEG study. NeuroImage. 2023 Jan 1;265:119802.

39. Truzzi A, Cusack R. The development of intrinsic timescales: A comparison between the neonate and adult brain. NeuroImage. 2023 Jul 15;275:120155.

40. Yates TS, Skalaban LJ, Ellis CT, Bracher AJ, Baldassano C, Turk-Browne NB. Neural event segmentation of continuous experience in human infants. Proceedings of the National Academy of Sciences. 2022 Oct 25;119(43):e2200257119.

41. Davis TS, Caston RM, Philip B, Charlebois CM, Anderson DN, Weaver KE, et al. LeGUI: A Fast and Accurate Graphical User Interface for Automated Detection and Anatomical Localization of Intracranial Electrodes. Front Neurosci [Internet]. 2021 Dec 9 [cited 2024 Nov 7];15. Available from: https://www.frontiersin.org/journals/neuroscience/articles/10.3389/fnins.2021.769872/full

42. Bokil H, Andrews P, Kulkarni JE, Mehta S, Mitra PP. Chronux: A platform for analyzing neural signals. Journal of Neuroscience Methods. 2010 Sep 30;192(1):146–51.

43. Donoghue T, Haller M, Peterson EJ, Varma P, Sebastian P, Gao R, et al. Parameterizing neural power spectra into periodic and aperiodic components. Nat Neurosci. 2020 Dec;23(12):1655–65.

44. Miller KJ, Sorensen LB, Ojemann JG, Den Nijs M. Power-Law Scaling in the Brain Surface Electric Potential. Sporns O, editor. PLoS Comput Biol. 2009 Dec 18;5(12):e1000609.

45. Gao R, Peterson EJ, Voytek B. Inferring synaptic excitation/inhibition balance from field potentials. NeuroImage. 2017 Sep 1;158:70–8.

46. Voytek B, Kramer MA, Case J, Lepage KQ, Tempesta ZR, Knight RT, et al. Age-Related Changes in 1/ f Neural Electrophysiological Noise. J Neurosci. 2015 Sep 23;35(38):13257–65.

47. He BJ, Zempel JM, Snyder AZ, Raichle ME. The Temporal Structures and Functional Significance of Scale-free Brain Activity. Neuron. 2010 May 13;66(3):353–69.

48. Linkenkaer-Hansen K, Nikouline VV, Palva JM, Ilmoniemi RJ. Long-Range Temporal Correlations and Scaling Behavior in Human Brain Oscillations. J Neurosci. 2001 Feb 15;21(4):1370–7.

49. Seabold S, Perktold J. Statsmodels: Econometric and Statistical Modeling with Python. In Austin, Texas; 2010 [cited 2025 Feb 3]. p. 92–6. Available from: https://doi.curvenote.com/10.25080/Majora-92bf1922-011

50. Benjamini Y, Hochberg Y. Controlling the False Discovery Rate: A Practical and Powerful Approach to Multiple Testing. Journal of the Royal Statistical Society: Series B (Methodological). 1995 Jan 1;57(1):289–300.

51. Johnson EL, Yin Q, O’Hara NB, Tang L, Jeong JW, Asano E, et al. Dissociable oscillatory theta signatures of memory formation in the developing brain. Current Biology. 2022 Apr;32(7):1457-1469.e4.

52. Groppe DM, Bickel S, Keller CJ, Jain SK, Hwang ST, Harden C, et al. Dominant frequencies of resting human brain activity as measured by the electrocorticogram. NeuroImage. 2013 Oct;79:223–33.

53. Frauscher B, Von Ellenrieder N, Zelmann R, Doležalová I, Minotti L, Olivier A, et al. Atlas of the normal intracranial electroencephalogram: neurophysiological awake activity in different cortical areas. Brain. 2018 Apr 1;141(4):1130–44.

54. Cross ZR, Gray SM, Dede AJO, Rivera YM, Yin Q, Vahidi P, et al. The development of aperiodic neural activity in the human brain [Internet]. bioRxiv; 2025 [cited 2025 Apr 23]. p. 2024.11.08.622714. Available from: https://www.biorxiv.org/content/10.1101/2024.11.08.622714v2

55. Baum GL, Flournoy JC, Glasser MF, Harms MP, Mair P, Sanders AFP, et al. Graded Variation in T1w/T2w Ratio during Adolescence: Measurement, Caveats, and Implications for Development of Cortical Myelin. J Neurosci. 2022 Jul 20;42(29):5681–94.

56. Burt JB, Demirtaş M, Eckner WJ, Navejar NM, Ji JL, Martin WJ, et al. Hierarchy of transcriptomic specialization across human cortex captured by structural neuroimaging topography. Nat Neurosci. 2018 Sep;21(9):1251–9.

57. Paquola C, Bethlehem RA, Seidlitz J, Wagstyl K, Romero-Garcia R, Whitaker KJ, et al. Shifts in myeloarchitecture characterise adolescent development of cortical gradients. Gold JI, Satterthwaite T, Mills K, Fulcher B, editors. eLife. 2019 Nov 14;8:e50482.

58. Grydeland H, Vértes PE, Váša F, Romero-Garcia R, Whitaker K, Alexander-Bloch AF, et al. Waves of Maturation and Senescence in Micro-structural MRI Markers of Human Cortical Myelination over the Lifespan. Cerebral Cortex. 2019 Mar 1;29(3):1369–81.

59. Baum GL, Cui Z, Roalf DR, Ciric R, Betzel RF, Larsen B, et al. Development of structure–function coupling in human brain networks during youth. Proceedings of the National Academy of Sciences. 2020 Jan 7;117(1):771–8.

60. Nenning KH, Xu T, Franco AR, Swallow KM, Tambini A, Margulies DS, et al. Omnipresence of the sensorimotor-association axis topography in the human connectome. Neuroimage. 2023 May 15;272:120059.

61. Wagstyl K, Ronan L, Goodyer IM, Fletcher PC. Cortical thickness gradients in structural hierarchies. NeuroImage. 2015 May;111:241–50.

62. Li G, Lin W, Gilmore JH, Shen D. Spatial Patterns, Longitudinal Development, and Hemispheric Asymmetries of Cortical Thickness in Infants from Birth to 2 Years of Age. J Neurosci. 2015 Jun 17;35(24):9150–62.

63. Vandekar SN, Shinohara RT, Raznahan A, Roalf DR, Ross M, DeLeo N, et al. Topologically Dissociable Patterns of Development of the Human Cerebral Cortex. J Neurosci. 2015 Jan 14;35(2):599–609.

64. Shaw P, Kabani NJ, Lerch JP, Eckstrand K, Lenroot R, Gogtay N, et al. Neurodevelopmental Trajectories of the Human Cerebral Cortex. J Neurosci. 2008 Apr 2;28(14):3586–94.

65. Hasson U. Uncovering a Timescale Hierarchy by Studying the Brain in a Natural Context. J Neurosci [Internet]. 2025 Mar 19 [cited 2025 Apr 18];45(12). Available from: https://www.jneurosci.org/content/45/12/e2368242025

66. Jensen O, Mazaheri A. Shaping Functional Architecture by Oscillatory Alpha Activity: Gating by Inhibition. Front Hum Neurosci [Internet]. 2010 Nov 4 [cited 2024 Oct 31];4. Available from: https://www.frontiersin.org/journals/human-neuroscience/articles/10.3389/fnhum.2010.00186/full

67. Samaha J, Iemi L, Haegens S, Busch NA. Spontaneous Brain Oscillations and Perceptual Decision-Making. Trends in Cognitive Sciences. 2020 Aug 1;24(8):639–53.

68. VanRullen R. Perceptual Cycles. Trends in Cognitive Sciences. 2016 Oct 1;20(10):723–35.

69. Samaha J, Postle BR. The Speed of Alpha-Band Oscillations Predicts the Temporal Resolution of Visual Perception. Current Biology. 2015 Nov 16;25(22):2985–90.

70. Torralba Cuello M, Drew A, Sabaté San José A, Morís Fernández L, Soto-Faraco S. Alpha fluctuations regulate the accrual of visual information to awareness. Cortex. 2022 Feb 1;147:58–71.

71. Bollimunta A, Chen Y, Schroeder CE, Ding M. Neuronal Mechanisms of Cortical Alpha Oscillations in Awake-Behaving Macaques. J Neurosci. 2008 Oct 1;28(40):9976–88.

72. Bollimunta A, Mo J, Schroeder CE, Ding M. Neuronal Mechanisms and Attentional Modulation of Corticothalamic Alpha Oscillations. J Neurosci. 2011 Mar 30;31(13):4935–43.

73. Raposo I, Fiebelkorn IC, Lin JJ, Parvizi J, Kastner S, Knight RT, et al. Human attention-guided visual perception is governed by rhythmic oscillations and aperiodic timescales. PLOS Biology. 2025 Jun 27;23(6):e3003232.

